# Prrx1-driven LINC complex disruption in vivo reduces osteoid deposition but not bone quality after voluntary wheel running

**DOI:** 10.1101/2023.09.22.559054

**Authors:** Scott Birks, Sean Howard, Christian S. Wright, Caroline O’Rourke, Elicza A. Day, Alexander J. Lamb, James R. Walsdorf, Anthony Lau, William R. Thompson, Gunes Uzer

**Author notes:** **Corresponding author:** Gunes Uzer PhD, Boise State University, Department of Mechanical & Biomedical Engineering, 1910 University Drive, MS-2085, Boise, ID 83725-2085.

## Abstract

The Linker of Nucleoskeleton and Cytoskeleton (LINC) complex serves to connect the nuclear envelope and the cytoskeleton, influencing cellular processes such as nuclear arrangement, architecture, and mechanotransduction. The role LINC plays in mechanotransduction pathways in bone progenitor cells has been well studied; however, the mechanisms by which LINC complexes govern *in vivo* bone formation remain less clear. To bridge this knowledge gap, we established a murine model disrupting LINC using transgenic Prx-Cre mice and floxed Tg(CAG-LacZ/EGFP-KASH2) mice. Prx-Cre mice express the Cre recombinase enzyme controlled by the paired-related homeobox gene-1 promoter (*Prrx1*), a pivotal regulator of skeletal development. Prx-Cre animals have been widely used in the bone field to target bone progenitor cells. Tg(CAG-LacZ/EGFP-KASH2) mice carry a lox-stop-lox flanked LacZ gene allowing for the overexpression of an EGFP-KASH2 fusion protein via cre recombinase mediated deletion of the LacZ cassette. This disrupts endogenous Nesprin-Sun binding in a dominant negative manner disconnecting nesprin from the nuclear envelope. By combining these lines, we generated a Prrx1(+) cell-specific LINC disruption model to study its impact on the developing skeleton and subsequently exercise-induced bone accrual. The findings presented here indicate Prx-driven LINC disruption (PDLD) cells exhibit no change in osteogenic and adipogenic potential compared to controls *in vitro* nor are there bone quality changes when compared to in sedentary animals at 8 weeks. While PDLD animals displayed increased voluntary running activity andPrrx1(+) cell-specific LINC disruption abolished the exercise-induced increases in osteoid volume and surface after a 6-week exercise intervention, no other changes in bone microarchitecture or mechanical properties were found.

## Introduction

Bone is a highly dynamic tissue influenced by both biochemical and mechanical stimuli to retain proper structure and function [1]. While the anabolic effects of mechanical challenge (i.e. weight-bearing exercise) on bone quality is well accepted, how Linker of Nucleoskeleton and Cytoskeleton (LINC) complexes may affect bone progenitor cell response to these signals remains underexplored.

LINC complexes serve as a physical link between the cytoskeleton and the nucleoskeleton, facilitating the transmission of mechanical forces and signals between the extracellular matrix and the nucleus [2]. LINC complexes are comprised of cytoskeletal nesprins (nuclear envelope spectrin repeats) and perinuclear Sun (Sad1/Unc84) proteins with the N-terminal calponin homology domains of nesprins binding to cytoskeletal F-actin microfilaments and the evolutionarily conserved C-terminal KASH (Klarsicht/Anc-1, Syne Homology) domains binding to Sun domains within the perinuclear space. Sun proteins directly engage with the intranuclear lamina, completing the connection from the cytoskeleton to the nucleus [3–5]. Beyond their structural role, LINC complexes have emerged as critical components in transducing mechanical signals within cells [6].

Earlier mesechmal stem cell (MSC) *in vitro* studies have effectively illustrated the dynamic involvement of LINC complexes within mechanotransduction pathways [6, 7]. For instance, disabling LINC complexes through RNA interference against SUN1 and SUN2 proteins, or transfecting cells with a dominant negative Nesprin KASH domain, leads to the suppression of focal adhesion kinase (FAK) and Akt phosphorylation[7]. This, in turn, circumvents the mechanical repression of adipogenesis when mesenchymal stromal cells are exposed to low-magnitude, high-frequency signals [6]. While the role of LINC complexes in mechanotransduction has been demonstrated *in vitro*, their role in orchestrating the mechanical regulation of bone in an *in vivo* remains largely uncharted. The current study aims to implement an *in vivo* strategy to investigate the ramifications of LINC complex disruption during mechanical loading, specifically within bone progenitor cells. This was achieved by creating a cre/lox Prrx1-driven LINC disruption (PDLD) murine model.

The paired-related homeobox gene-1 (*Prrx1*) is widely used to target bone progenitor cells as it is expressed in early limb bud mesenchyme, playing a crucial role in regulating skeletal development, and is found in adult undifferentiated mesenchyme and periosteum [8–11]. Thus, using transgenic Prrx1-cre mice we are able to target skeletal tissues throughout development and into adulthood. LINC disruption in Prx(+) cells was then achieved by crossing these mice with floxed mice created by Razafsky and Hodzic that overexpress an EGFP-KASH2 fusion protein upon Cre-recombination [12, 13]. This fusion protein binds to perinuclear Sun proteins in a dominant negative manner, causing displacement of cytoplasmic nesprins into the endoplasmic reticulum and thus, disrupting LINC connectivity between the nucleus and cytoskeleton.

While there are several modalities for mechanical stimulation *in vivo*, exercise is considered a potent modality that can trigger adaptive responses in bone tissue, enhancing bone density and strength [14]. A variety of exercise models including forced/trained treadmill running, voluntary/forced wheel running, swimming, and resistance training (loaded/unloaded ladder climbing) have been utilized in rodent studies, all of which have been shown to improve bone mass, bone strength, and bone mineral density [15] [16]. We chose voluntary wheel running for the present study to reduce the amount of handling/environmental stress on the animals.

To interrogate the effect of disabling LINC complex function in Prx positive bone cell progenitors, we created a LINC disruption model under the Prx-Cre driver and measured primary cell differentiation potential *in vitro*, bone microarchitecture via micro-computed tomography (micro-CT) and mechanical properties via 3-point bending at an 8-week baseline. We then subjected a cohort of animals to either a running or non-running intervention starting at 8-weeks of age and lasting for 6 weeks with a 1-week acclimation period. Micro-CT and biomechanical properties were also measured post-exercise to assess any effects of LINC disruption on exercise induced bone accrual.

## Materials and Methods

### Animals

Mouse strains included B6.Cg-Tg(Prrx1-cre))1Cjt/J aka *Prx-Cre* (The Jackson Laboratory, Stock Number: 005584)[8], Tg(CAG-LacZ/EGFP-KASH2) aka *KASH2* (Hodzic, Wash U)[12], and B6.Cg-Gt(ROSA)26Sor^tm14(CAG-tdTomato)Hze^/J aka *Ai14* (generously provided by Dr. Richard Beard)[17].

Hemizygous *Prx-Cre* mice were crossed with floxed *KASH2* mice to generate Prrx1-driven LINC disrupted (PDLD) murine model. LINC disruption mechanism described by Razafsky and Hodzic [12, 13]. Briefly, upon cre recombination, a LacZ containing lox-stop-lox cassette upstream of an eGFP-KASH2 ORF is excised allowing for the overexpression an eGFP-KASH2 fusion protein which saturates available Sun/KASH binding in a dominant-negative manner. Genotyping was performed on tissue biopsies via real-time PCR probing for LacZ and Cre (Transnetyx, Cordova, TN) to determine experimental animals and controls. Cre(+)/LacZ(+) animals were considered experimental and all other genotypes were considered controls. Animals were housed in individually ventilated cages under controlled conditions until placement in conventional cages for the duration of the exercise intervention. *Ad libitum* access to food and water allowed.

Additionally, *Prx-Cre* mice were crossed with Ai14 RFP reporter mice to assess Prx-Cre activity in skeletal tissue and bone marrow cells. All procedures approved by Boise State University Institutional Animal Care and Use Committee. During sacrifice animals were first anesthetized via Isoflurane inhalation followed by decapitation in this way efforts to alleviate suffering. Except sacrifice animals were subjected to running procedures which did not cause any distress or did not require the use of pain-relieving drugs.

### Bone Marrow Cell Isolation, Expansion, and Differentiation

Bone marrow cells were isolated from tibiae and femora via centrifugation as previously described [18]. Briefly, tibia and femur were dissected, cleared of all soft tissue, and centrifuged at 12,000xg to remove the bone marrow. This bone marrow pellet was then resuspended in media and plated in supplemented growth media (alpha-MEM +20% FBS, +100U/mL penicillin, +100µg/mL streptomycin). Cells were washed with PBS after 48 hours to remove non-adherent cells and allowed to proliferate to 50% confluency. At this confluency, cells were lifted, split onto two separate plates to produce enough cells for assays, and allowed to proliferate to ∼80% confluency. Cells were then lifted and plated for differentiation assays.

For osteogenic assays (n=4 replicates/group), cells were plated in triplicate and seeded at 200K per well on 6-well plates in normal growth media for 24 hours prior to switching to osteogenic media (alpha-MEM with 20% FBS, 100U/mL penicillin, 100µg/mL streptomycin, 10mM beta-glycerophosphate, 50µg/µL ascorbic acid) for 14 or 21 days.

For adipogenic assays (n=4 replicates/group), cells were plated in triplicate and seeded at 150K per well on 6-well plates in normal growth media for 24 hours prior to switching to adipogenic media (alpha-MEM with 10% FBS, 100U/mL penicillin, 100µg/mL streptomycin, 5µg/mL insulin, 0.1µM dexamethasone, 50µM indomethacin) for 7 days.

### Immunofluorescence staining and microscopy

Nesprin-1, Nesprin-2, Sun-1, and Sun-2 staining was performed on bone marrow cultures from 12-week Prx-Cre(+)/eGFP-KASH2(+) and aged-matched control mice (n=2/group). After isolation and culture as described above, cells were rinsed with PBS, fixed in 4% paraformaldehyde, permeabilized with 0.3% Triton buffer and blocked with 5% goat serum (Jackson ImmunoResearch 005-000-001) prior to being tagged with a primary antibody.. After primary antibody tagging, cells were incubated with an anti-rabbit secondary antibody (1:300 AlexaFluor 594nm goat anti-rabbit, Invitrogen A11037), nuclei were stained with Hoechst 33342 (Invitrogen™ R37605) and visualized via fluorescent microscopy (Zeiss LSM 900). No additional staining was needed to visualize endogenous eGFP. Following primary antibodies were used, Sun-1 (Sigma Aldrich, HPA008346), Sun-2 (Abcam, ab87036), Nesprin-2 (ImmuQuest, IQ565), Nesprin-1 (Millipore Sigma, MABT843).

### Oil Red O and Alizarin Red Staining

After 7 days in adipogenic media, cultures were stained against Oil Red O as described previously [19]. Briefly, cells were rinsed with PBS, fixed in 10% neutral buffered formalin for 30 mins at room temperature, and washed with deionized water. Cultures were then incubated in 100% propylene glycol, stained with 0.5% Oil Red O in propylene glycol, and finally incubated in 85% propylene glycol. Unbound dye was removed by rinses with tap water until clear and images were taken. Oil Red O stain was then eluted with 100% propan-2-ol and optical absorbance values were read on a microplate spectrophotometer (ThermoFisher Multiskan GO) at 510nm.

After 14 days in osteogenic media, cells were rinsed in PBS, fixed with 70% ethanol, and stained with 40mM Alizarin Red solution for visualization of calcium deposits. The stain was then eluted using 10% w/v cetylpyridium chloride (CPC) and diluted 4-fold to avoid absorbance saturation. Absorbance values were read on a microplate spectrophotometer (ThermoFisher Multiskan GO) at 562nm.

### Western Blot

Lysates were prepared using radio immunoprecipitation assay (RIPA) lysis buffer (150mM NaCl, 50mM Tris HCl, 1mM EDTA, 0.24% sodium deoxycholate,1% Igepal, pH 7.5) to protect the samples from protein degradation NaF (25mM), Na3VO4 (2mM), aprotinin, leupeptin, pepstatin, and phenylmethylsulfonylfluoride (PMSF) were added to the lysis buffer. Western protein amounts were normalized to 15μg through BCA Protein Assay (Thermo Scientific, #23225). Lysates (20μg) were separated on 10% poly-acrylamide gels and transferred to polyvinylidene difluoride (PVDF) membranes. Membranes were blocked with milk (5%, w/v) diluted in Tris-buffered saline containing Tween20 (TBS-T, 0.05%). Blots were then incubated overnight at 4°C with appropriate primary antibodies. Following primary antibody incubation, blots were washed and incubated with horseradish peroxidase-conjugated secondary antibody diluted at 1: 5,000 (Cell Signaling) at RT for 1h in 5% milk in TBST-T. Chemiluminescence was detected with ECL plus (Amersham Biosciences, Piscataway, NJ). Following antibodies were used, Sun-1 (Sigma Aldrich, HPA008346), Sun-2 (Abcam, ab87036, eGFP (abcam, ab184601)

### Exercise Regimen

Eight-week old male PDLD or control mice were randomly assigned to a voluntary wheel running exercise intervention, upright running wheel (Kaytee Silent Spinner Exercise Wheel), or a non-running control group without access to a running wheel (n=10-12 mice/group). Animals were allotted a one-week acclimation period in these new cages followed by a six-week tracked exercise intervention. All mice were individually housed and running metrics including elapsed time, total distance, average speed, and maximum speed were measured with a cyclocomputer (CatEye Velo 7) and recorded daily. Ad libitum access to food and water was allowed. Body mass and food intake was also tracked weekly.

### Micro-computed tomography

Proximal tibiae (n=10-12 mice/group) stored frozen in PBS soaked gauze prior to scanning and were scanned immersed in PBS using an x-ray microtomograph (SkyScan 1172F). Settings as follows: power of 55kV/181mA, a voxel size of 10.33µm^3^, and 230ms integration time. The volume of interest for trabecular quantitative analysis started 10 slices distal to the proximal physis extending 1 mm. The cortical volume of interest started 2.15mm distal to the physis, extending another 0.5mm. Image reconstruction was performed with NRecon software (Bruker). After any necessary image reorientation, quantitative analysis was performed with CTAn (Bruker) and 3D image rendering was performed with CTVol. Reported variables and nomenclature used according to ASBMR guidelines [20].

### Three-point Bend Mechanical Testing

Bone strength was evaluated using 3-point bending on the tibias [21–23]. Tibias were rehydrated in PBS for at least 24 hours prior to performing biomechanical testing. Biomechanical testing was performed using a 3-point bending test fixture with an Instron Universal Testing System model 5967. Due to the asymmetric nature of the tibia cross section, the biomechanical test was optimized to place the most uniform cross section of the tibia between the spans, which was set to 7.00 mm. This resulted in a region just below the trochanter and the distal end of the tibia. A downward displacement of 0.02mm/s was applied until failure. Force and deflection data was collected and analyzed in using MATLAB in order to determine the mechanical properties of the bone(i.e. linear flexural stiffness, maximum yield force, and maximum force) according to previously published methods [24, 25].

### Histology

Tibias from 6-week voluntary running (n=8/grp) were MMA-embedded, thin sections were deplasticized in acetone and stained via a modification of the von Kossa / MacNeal’s (VKM) Tetrachrome protocol.[26] For VKM-stained slides, mineralized bone was stained using the Von Kossa silver method and the unmineralized tissue was counter-stained with MacNeal’s tetrachrome. Histomorphometric analysis was performed using the OsteoMeasure high resolution digital video system (OsteoMetrics Inc., Decatur, GA). Standard nomenclature was used according to the Histomorphometry Nomenclature Committee of the American Society of Bone and Mineral Research.[27] Measurements included: osteoid volume per bone volume (OV/BV), osteoid surface per bone surface (OS/BS), osteoblast surface per bone surface (Ob.S./BS.), number of osteoblasts per bone perimeter (N.Ob./B.Pm.), and number of osteoblasts per osteoblast perimeter (N.Ob./Ob.Pm.).

### RNA-sequencing

Total RNA was extracted using RNAeasy (Qiagen) for three samples per group. Total RNA samples were sent to Novogene for mRNA sequencing and analysis. Briefly, index of the reference genome was built using Hisat2 v2.0.5 and paired-end clean 2 reads were aligned to the reference genome using Hisat2 v2.0.5. featureCounts v1.5.0-p3 was used to count the reads numbers mapped to each gene. And then FPKM of each gene was calculated based on the length of the gene and reads count mapped to this gene. FPKM, expected number of Fragments Per Kilobase of transcript sequence per Millions base pairs sequenced. Differential expression analysis was performed using the DESeq2 R package (1.20.0). DESeq2 provides statistical routines for determining differential expression in digital gene expression data using a model based on the negative binomial distribution. The resulting P-values were adjusted using the Benjamini and Hochberg’s approach for controlling the false discovery rate. Genes with an adjusted P-≤0.05 found by DESeq2 were assigned as differentially expressed. Genes with significant differential gene expression were further analyzed with DAVID[28] for pathway analysis. Pathways with a Benjamini ≤0.05 were selected, results were presented in Tables S1 and S2.

### Statistical Analysis

Results are expressed as mean ± standard deviation. Normality was tested via Shapiro–Wilk test. Significance was determined by 2-tailed unpaired student’s t-test or 2-way ANOVA with Tukey post-hoc test (Graphpad Prism version 10.2.0).

## Results

### Expression of EGFP-KASH2 fusion protein physically disrupts LINC complex in Prrx1 positive cells

We crossed Prx-Cre(+) mice with the Ai14 RFP reporter line to visualize Cre activity in skeletal tissues and bone marrow cells. P7 pups were placed in an *in vivo* fluorescent scanner (IVIS® Spectrum) and notable red fluorescent in the appendicular skeleton and calvaria was visible (Figure 1A), consistent with previous reports [29]. This confirms Cre recombinase activity is present in Prrx1(+) tissues. We then isolated bone marrow from these animals at 12 weeks and successfully visualized RFP containing cells within the bone marrow (Figure 1B) providing evidence for the presence of Prrx1-Cre activity in bone marrow.

**Figure 1.**
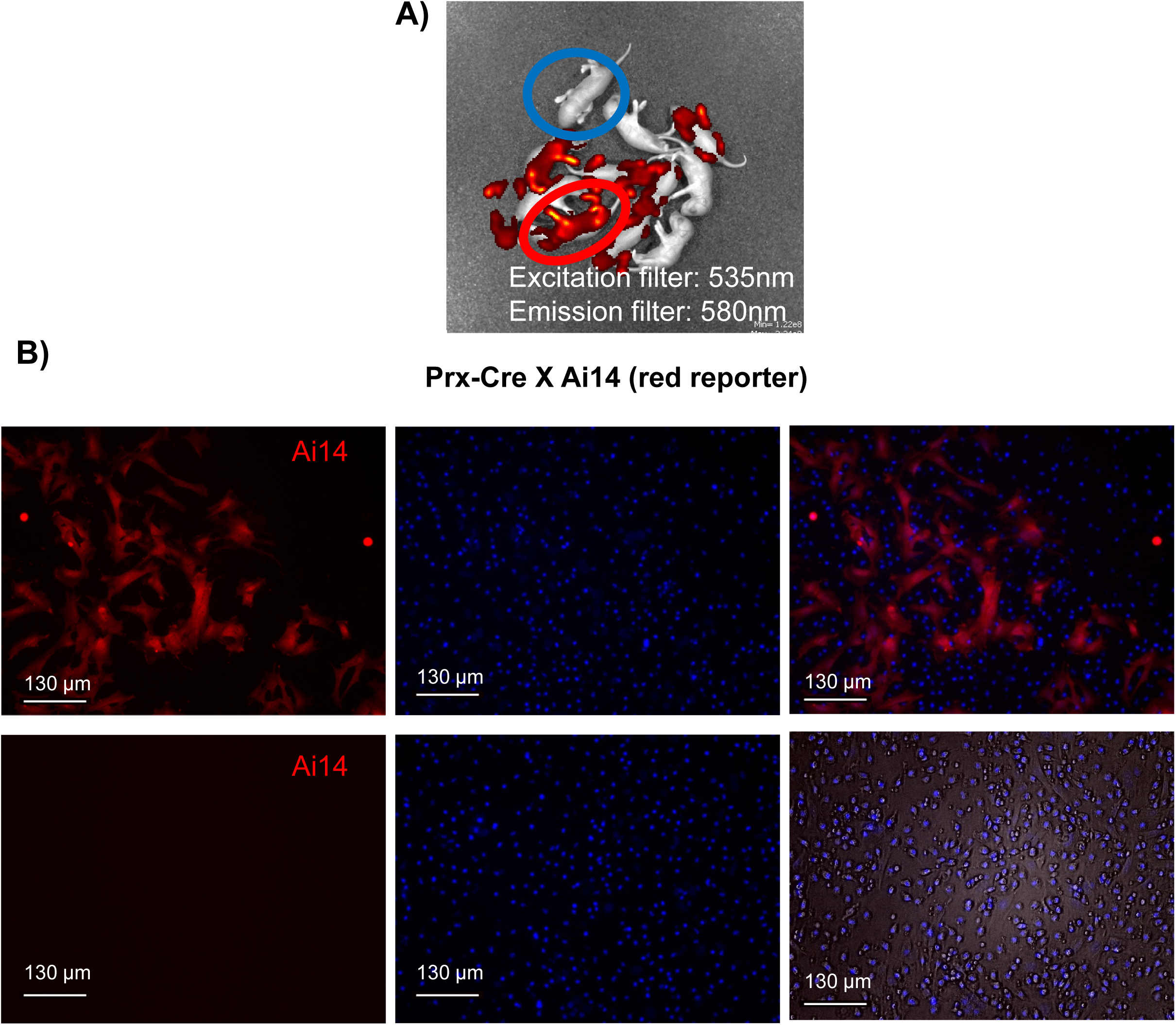
Prx-Cre activity confirmed in bone marrow using Ai14 reporter animals. **A)** IVIS® Spectrum *in vivo* fluorescent scan of P7 pups from Prrx1-Cre/Ai14 cross indicating skeletal Cre activity in Prx-Cre(+) mice (red) vs. no Cre activity in Prx-Cre(-) littermates (blue). **B)** Representative image of bone marrow aspirates from 8-week male Prx-Cre(+)/Ai14 mice indicating the presence of Prx-Cre(+) cells within the bone marrow space.

A Prx-driven LINC disruption (PDLD) model was then generated by crossing hemizygous Prx-Cre(+) mice with floxed Tg(CAG-LacZ/EGFP-KASH2) mice. The mechanism for LINC disruption in this model relies on Cre recombinase mediated excision of the lox-stop-lox LacZ cassette in the *KASH2* mice allowing for the overexpression of an eGFP-KASH2 fusion protein. This fusion protein then binds to perinuclear Sun proteins, displacing nesprins from the nuclear envelope.

To confirm this model disrupts LINC function via displacement of LINC complex elements nesprin1, nesprin-2, sun-1 and sun-2, we flushed bone marrow from 12-week old PDLD and control animals (N=4/group) to check for cells expressing endogenous eGFP localized to the nuclear envelope, indicating the overexpression of the eGFP-KASH2 protein. We additionally immunostained these cells with an anti-nesprin1, nesprin-2 (Figure2), sun-1 and sun-2 (Figure 3) antibodies and secondary fluorophore to confirm their detachment or decreased expression from the nuclear envelope. As seen in Figure 2 and 3, there is a clear nuclear halo of eGFP fluorescence in PDLD mice compared to no eGFP visualized in controls, we also confirmed the eGFP expression via bone marrow flushing followed by western blotting agans eGFP (Fig.S1A).

**Figure 2.**
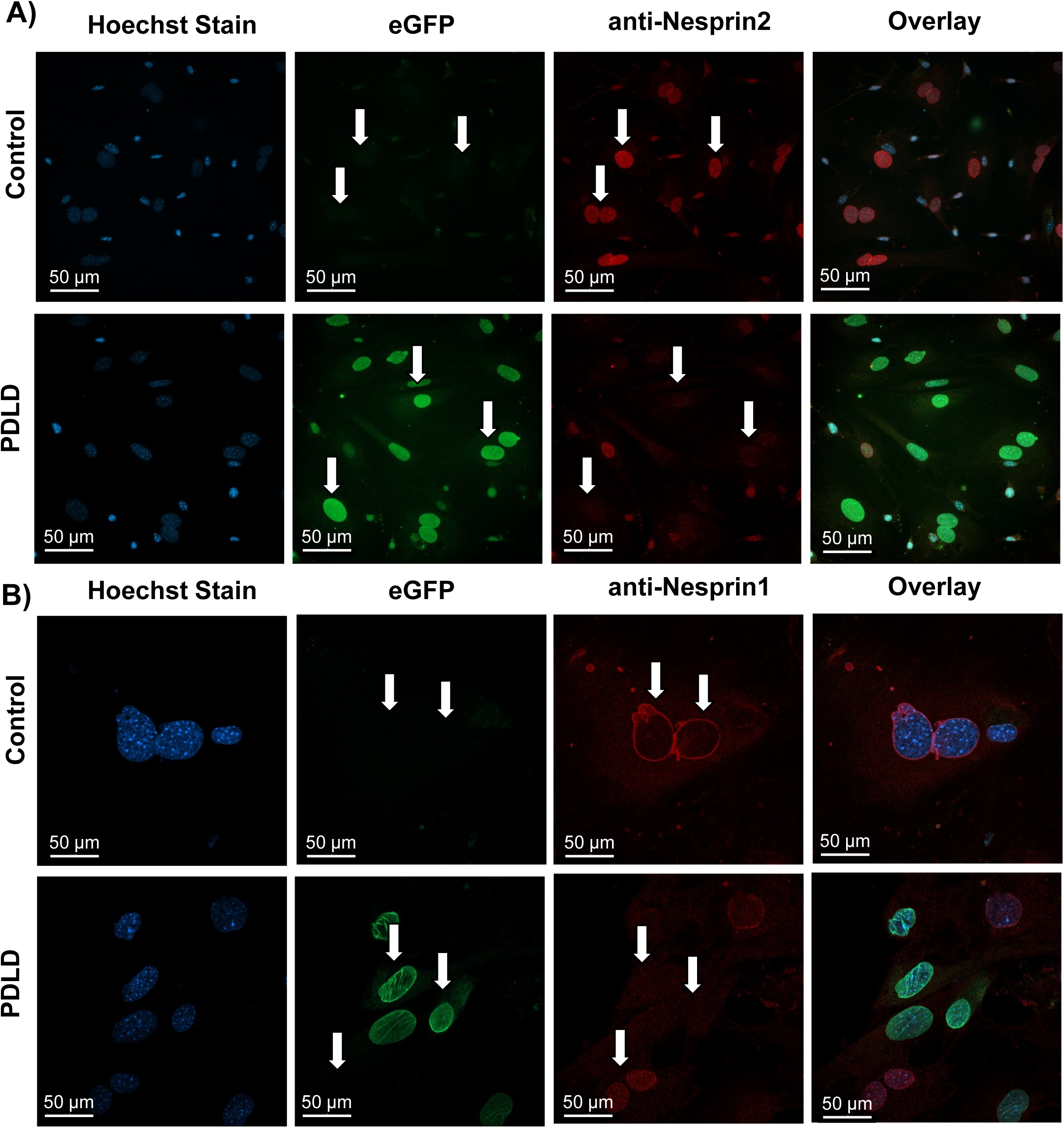
Prrx1-driven LINC disruption (PDLD) model characterization. **A)** Bone marrow aspirates from 12-week male Prrx1-driven LINC disrupted (PDLD) and control littermates. Confirmation of eGFP expression localized to the nuclear envelope with nesprin displacement compared to no eGFP visualization and intact nesprin localization to the nuclear envelope in controls. Cells were manually counted and analyzed for nuclear localization of eGFP and/or anti-nesprin 2 red fluorescence. In PDLD cultures: 72.4% of cells had green, but not red nuclear localization; 17.7% has red, but no green nuclear localization; and 9.9% had both (n=141 cells from 12 images). In control cultures: out of 12 images from two separate animals, there was no appreciable green fluorescence from 121 cells with red fluorescent nuclear localization. **B)** Bone marrow aspirates from 12-week male Prrx1-driven LINC disrupted (PDLD) and control littermates. Confirmation of eGFP expression localized to the nuclear envelope with nesprin displacement compared to no eGFP visualization and intact nesprin localization to the nuclear envelope in controls. Cells were manually counted and analyzed for nuclear localization of eGFP and/or anti-nesprin 1 red fluorescence.

**Figure 3.**
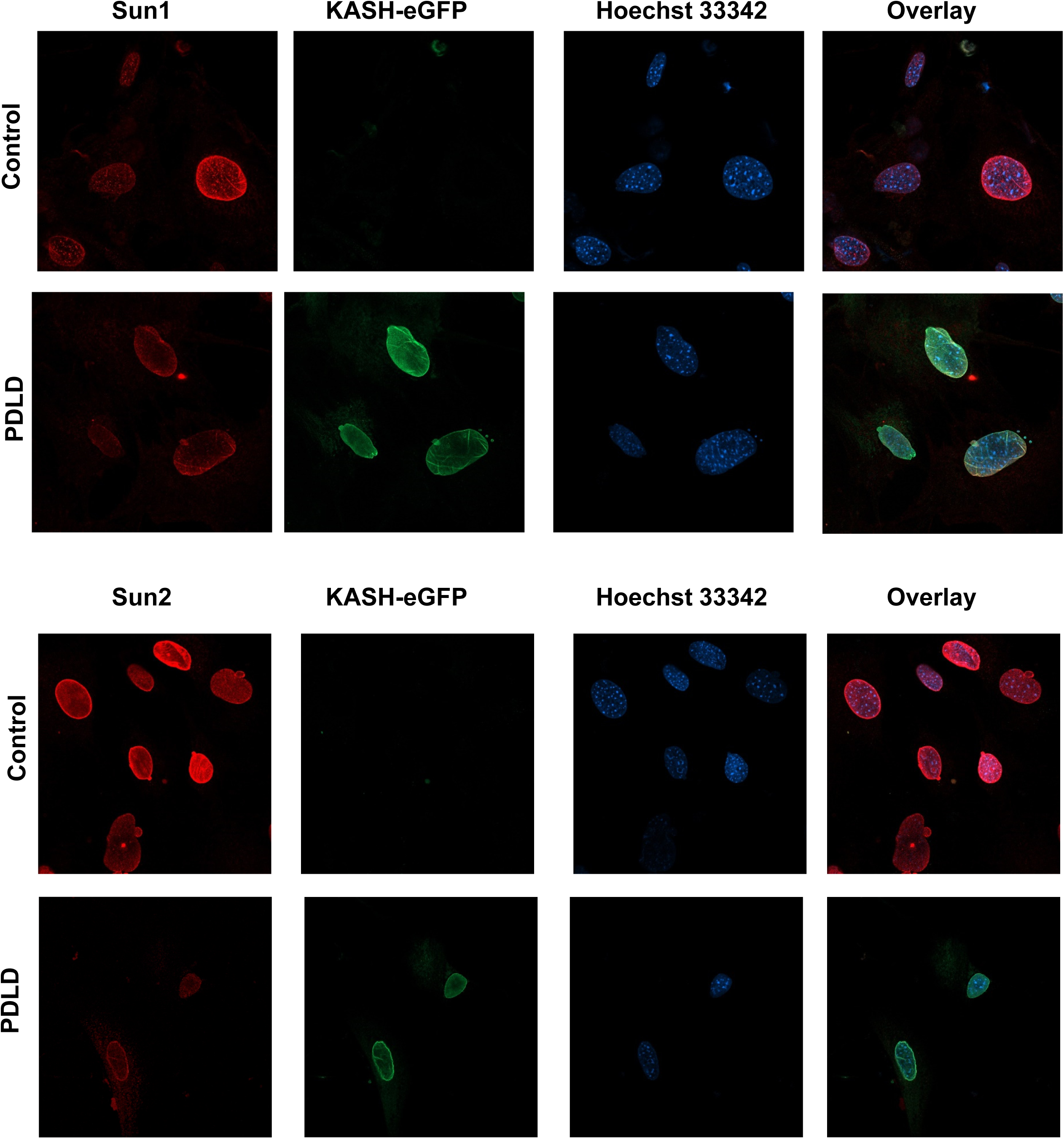
Prrx1-driven LINC disruption (PDLD) model characterization. **B)** Representative image of bone marrow aspirates from 8-week male Prx-Cre(+)/Ai14 mice indicating the presence of Prx-Cre(+) cells within the bone marrow space. **C)** Bone marrow aspirates from 12-week male Prrx1-driven LINC disrupted (PDLD) and control littermates. Confirmation of eGFP expression localized to the nuclear envelope with nesprin displacement compared to no eGFP visualization and intact nesprin localization to the nuclear envelope in controls.

Shown in Figure 2, nesprin-1 and nesprin-2 stainings do not show a clear nuclear localization pattern (nuclear halo) in PDLD mice compared to the clear nuclear positioning seen in controls indicating successful displacement of nesprins from the nuclear envelope where eGFP is present. Shown in Figure 3 and Fig. S1B, sun1 and sun 2 show a slight decrease in PDLS animas. These data provide evidence of successful LINC disruption in cells expressing eGFP.

### Prrx1-driven LINC disruption does not change adipogenic and osteogenic differentiation potential from LINC-intact controls

To determine the differentiation potential of LINC disrupted cells, bone marrow was flushed from tibia and femur of 12-week male PDLD and negative control mice (n=4) and plated for osteogenic/adipogenic differentiation assays (Figure 4). After 7 days, adipogenic cultures were stained with Oil Red O and eluted stain was quantified via spectrophotometry at 510nm. After 14 days, osteogenic cultures were stained with Alizarin Red and eluted stain was quantified via spectrophotometry at 562nm.

**Figure 4.**
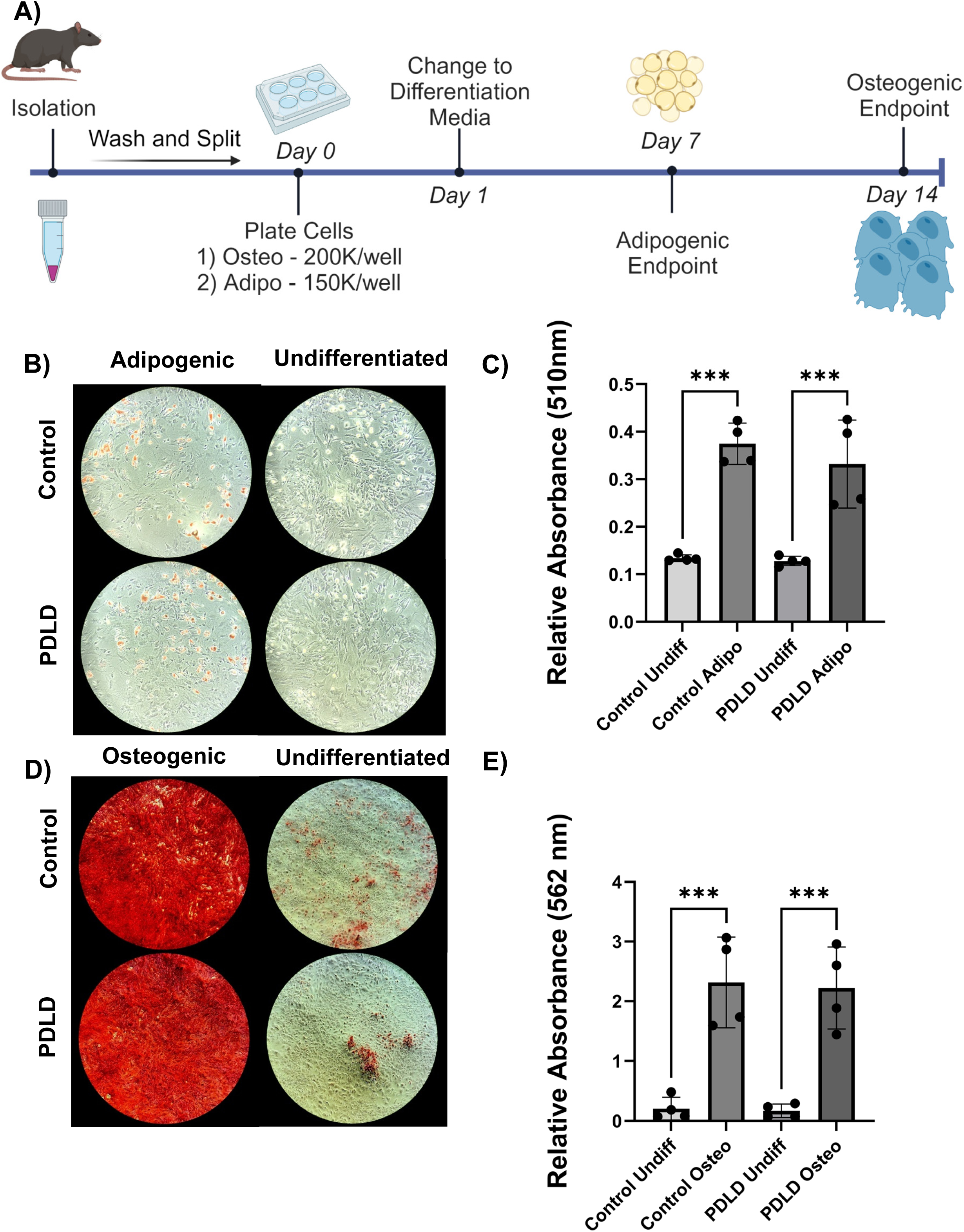
Bone marrow aspirates from PDLD mice do not display differences in osteogenic and adipogenic differentiation potential compared to LINC intact controls. **A)** Schematic for differentiation assay timeline as described in Methods **B)** 4x images of Oil Red O staining after 7 days in adipogenic or normal growth media for 12 week male ODLD mice vs LINC intact controls (n=4/grp). **C)** Absorbance values for eluted Oil Red O stain read at 510nm. **D)** 4x images of Alizarin Red staining after 21 days in osteogenic or normal growth media. **E)** Absorbance values for eluted Alizarin Red stain read at 562nm. Results presented as mean +/-SD. Normality confirmed with Shapiro-Wilk test. Significance determined via ordinary one-way ANOVA. *p<0.05, **p<0.01, ***p<0.001

Relative absorbance values of Oil Red O dye from PDLD adipogenic cultures was 24.8% lower than adipogenic LINC-intact controls (p=0.15) and 120.4% greater than undifferentiated controls (p<0.05). Relative absorbance values of Alizarin Red stain from PDLD osteogenic cultures was 5.0% less than that of LINC intact osteogenic controls (p=0.8), and 531.4% greater than undifferentiated LINC-intact controls (p<0.01). This suggests a trend towards diminished adipogenic differentiated potential in PDLD cells compared to LINC-intact controls, but no appreciable difference in osteogenic potential.

To exclude the possibility of these cells not surviving the marrow extraction we first confirmed that eGFP-KASH expressing LINC-deficient cells were present in our cultures after marrow extraction. Shortly, we harvested cells from female Prx-Cre(+)/Ai14(+)/eGFP-KASH2(+) animals that express eGFP and tdTomato in Prx(+) cells. Cells were visualized for green and red fluorescence. Average cell counts from 24 representative images with 683 total cells yielded 98.07 ± 7.2% red cells and 86.3 ± 14.52 % green cells indicating that eGFP-KASH2 expressing cells were present in our cultures (Fig. S2 A/B). Next to confirm participation of LINC-deficient cells in osteogenic nodules we have differentiated these cultures into osteogenic lineage and stained with calcein blue. Our results in Fig.S2 C confirms that Prx-Cre(+)/eGFP-KASH2(+) cells were present within calcein blue stained nodules. Finally, it is worth noting that the duration of *in vitro* studies involving KASH overexpression is relatively short (1-2 days to 1 week), while the animals in this study express the dominant negative KASH proteins chronically from conception. To test the possible effect of acute versus chronic KASH protein expression, we utilized transient transfection with a previously published dominant negative form of the nesprin-1 KASH domain (DNK) [30, 31]. RNA-seq and qPCR results from transient DNK transfection show decreased osteogenic gene transcripts *in vitro* when compared to empty mCherry plasmid transfected controls (Fig.S3). Pathway analysis were provided in Tables S1 and S2 indicates decreased collagen production was decreased in DNK expressing groups.

### Bone microarchitecture unchanged in PDLD mice compared to LINC-intact controls at 8-week baseline

Prior to starting a cohort on the exercise intervention, we established baseline bone microarchitecture measurements via micro-computed tomography (micro-CT) using an 8-week cohort of mice covering all four possible genotypes (n=3/grp): Prx-Cre(+)/EGFP-KASH2(+), Prx-Cre(+)/EGFP-KASH2(-), Prx-Cre(-)/EGFP-KASH2(+), Prx-Cre(-)/EGFP-KASH2(-). There was no significant difference found between genotypes (Fig. S4), so Prx-Cre(+)/EGFP-KASH2(+) mice remained our experimental model while all other genotypes were indiscriminately used as controls.

Each mouse from this cohort was weighed and both tibiae and femora were harvested from Prx-Cre(+)/EGFP-KASH2(+) and control (n=10/group). Each tibia was cleared of soft tissue for micro-CT analysis. We noted no change in body weight in PDLD mice vs. control recorded at 21.59 ± 1.7 g and 21.20 ± 0.77 g (p=0.68) respectively. A trend towards decreased bone microarchitecture with a 5.6% decrease in BV/TV (p=0.30), 5.3% decrease in trabecular thickness (p=0.19), and 6.10% decrease in cortical bone area (p=0.12) in PDLD mice was observed between control and PDLD mice (Figure 5).

**Figure 5.**
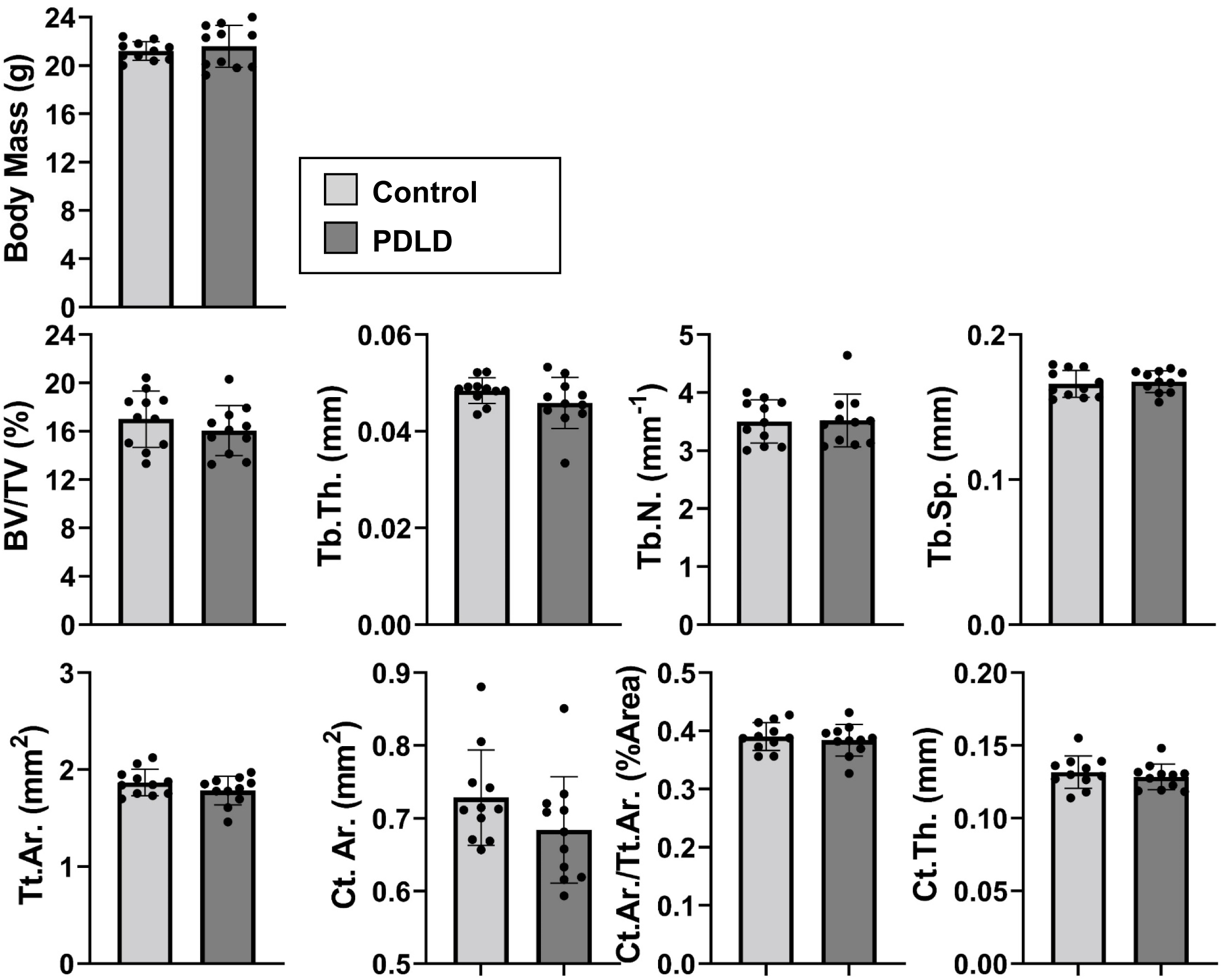
Bone microarchitecture and body mass not significantly different between PDLD and control mice at 8-week baseline. Trabecular measurements include bone volume fraction (BV/TV), trabecular thickness (Tb.Th.), trabecular spacing (Tb.Sp.), and trabecular number (Tb.N). Cortical measurements include total cross-sectional area inside the periosteal envelope (Tt.Ar.), cortical bone area (Ct.Ar.), cortical area fraction (Ct.Ar./Tt.Ar.), and average cortical thickness (Ct.Th.) Significance determined via non-parametric Student’s t-test. *p<0.05, **p<0.01, ***p<0.001

### Prrx1-driven LINC disrupted mice run longer, farther, and faster than controls

A second cohort of 8-week male mice were given voluntary access to a running wheel (or no wheel and allowed a one-week acclimation period before beginning the six-week tracked exercise intervention. Tracked running metrics including elapsed time, average distance, average velocity, and max speed were recorded daily (Figure 6). Our data indicate that PDLD mice ran significantly longer, farther, and faster than their LINC intact counterparts with an average run time of 169.2 ± 6.8 mins compared to 146.6 ± 9.4 mins (p<0.001), an average distance of 2.35 ± 0.16 km compared to 1.82 ± 0.1 km (p<0.0001), and an average velocity was 0.53 ± 0.05 km/hr for PDLD mice and 0.48 ± 0.02 km/hr for controls (p<0.05).

**Figure 6.**
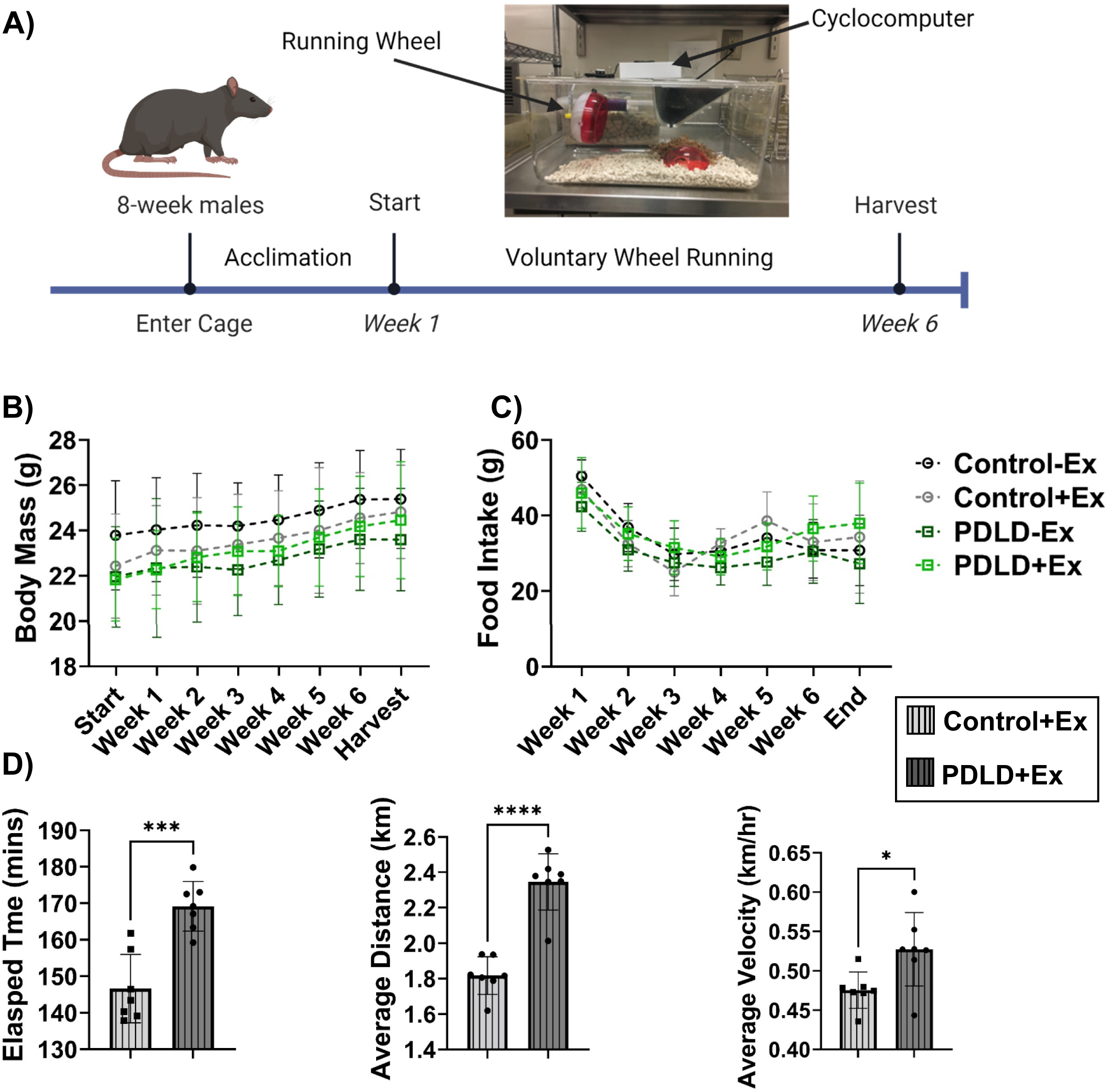
PDLD mice run significantly longer, farther, and faster than controls. **A)** Exercise regimen timeline schematic **B)** Body mass steadily increased throughout exercise intervention with no significant difference between groups. **C)** No significant difference in food intake between groups. **D)** PDLD mice ran longer, farther, and faster than LINC intact controls. Running metrics tracked by cyclometer measuring elapsed time, average distance and average velocity. Significance determined via 2-way ANOVA for body weight and food intake and unpaired Student’s t-test for elapsed time, average distance, and average velocity. *p<0.05, **p<0.01, ***p<0.001

### LINC disruption does not significantly affect bone microarchitecture or mechanical properties in PDLD mice subjected to six-week running protocol but decreases exercise-induced osteoid deposition

Bone microarchitecture was measured via micro-CT to assess any effects of LINC disruption on exercise-induced bone accrual in the proximal tibial metaphysis and diaphysis for trabecular and cortical parameters, respectively. There was no notable difference in trabecular bone fraction (BV/TV), trabecular thickness (Tb.Th.), trabecular separation (Tb.Sp.), trabecular number (Tb.N), total cortical tissue area (Tt.Ar.), or cortical thickness (Ct.Th.) between any of the groups. There was, however, a 10.5% decrease (p<0.05) in cortical bone area (Ct.Ar.) in PDLD+Ex compared to sedentary LINC-intact controls and a 6.15% decrease (p<0.05) in cortical area fraction (Ct.Ar./Tt.Ar.) in PDLD+Ex compared to PDLD-Ex. No significant changes in measured bone mechanical properties were noted (Figure 7). Representative micro-CT images for trabecular and cortical architecture of control and PDLD animals with and without exercise were presented in Figure S6.

**Figure 7.**
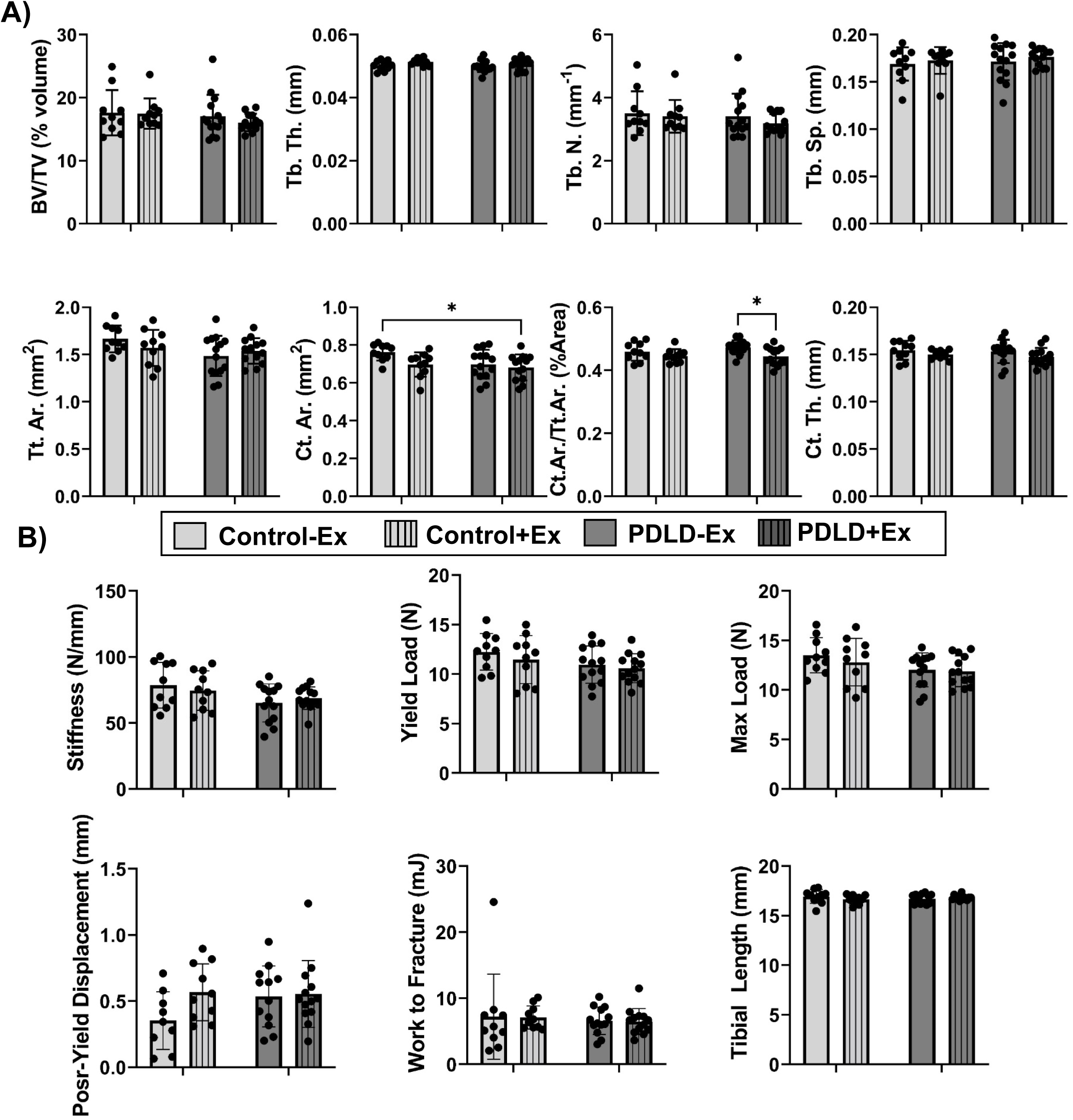
Trabecular and cortical microarchitecture of proximal tibiae unaffected by LINC disruption after 6-week voluntary wheel running. Trabecular measurements include bone volume fraction (BV/TV), trabecular thickness (Tb.Th.), trabecular spacing (Tb.Sp.), and trabecular number (Tb.N). Cortical measurements include total cross-sectional area inside the periosteal envelope (Tt.Ar.), cortical bone area (Ct.Ar.), cortical area fraction (Ct.Ar./Tt.Ar.), and average cortical thickness (Ct.Th.) n=10-12 mice/group. Representative images provided in Figure S6. Significance determined two-way ANOVA with Tukey post-hoc test ‘*p<0.05, **p<0.01, ***p<0.001

Tibias from 6-week voluntary running (n=8/grp) were MMA-embedded and stained via a modification of the von Kossa / MacNeal’s (VKM) Tetrachrome protocol.[26] Shown in Fig.8, analysis indicate that voluntary running exercise significantly increases osteoid volume (OV/BV, 252%, p<0.01) and osteoid surface (OS/BS, 235%, p<0.05) in control animals, suggesting that exercise increases organic portion of the bone matrix that forms prior to the maturation. In PDLD mice however this response to voluntary running exercise was completely abolished, decreasing averaged osteoid volume by 17-fold (OV/BV, p<0.0001) and that osteoid surface by 9.7-fold in exercised PDLD animals when compared to exercised control animals. Non-exercised osteoid measures of control animals were not significantly different from non-exercised PDLD mice. While no exercise effect was observed on osteoblast measures under exercise and PDLD conditions, exercised PDLD mice showed a modest decrease of Osteoblasts surface per bone surface (Ob.S./BS) when compared to both non exercised (42%, p<0.0.5) and exercised (50%, p<0.0.5) control mice. Similarly, osteoblast number per bone perimeter (N.Ob./Ob.Pm) of PDLD exercise mice showed a small decrease (60%, p<0.05) when compared to non-exercised control. No changes in of osteoblasts per osteoblast perimeter (N.Ob./Ob.Pm.) were found.

**Figure 8.**
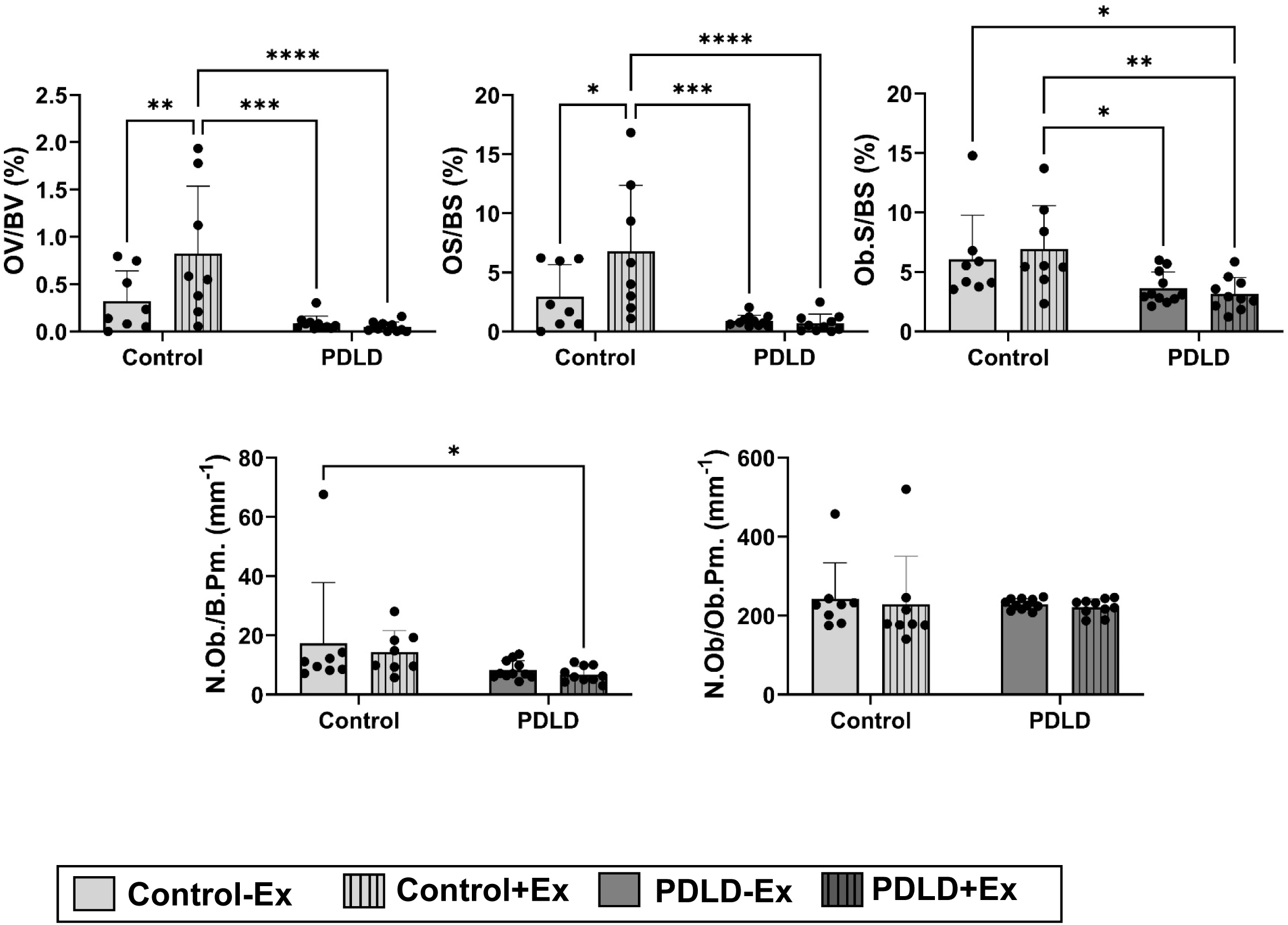
Von kossa staining for proximal tibiae section in LINC-disrupted and LINC-intact animals after 6-week voluntary wheel running. Measurements included: osteoid volume per bone volume (OV/BV), osteoid surface per bone surface (OS/BS), osteoblast surface per bone surface (Ob.S./BS.), number of osteoblasts per bone perimeter (N.Ob./B.Pm.), and number of osteoblasts per osteoblast perimeter (N.Ob./Ob.Pm.). n=8 animal per group.*p<0.05, **p<0.01, ***p<0.001

## Discussion

While the *in vitro* role of LINC complex in mechanotransduction pathways in bone progenitor cells has been relatively well studied [32–45], its function *in vivo* remains underexplored. Here we sought to elucidate if LINC complex disruption in *Prx(*+) cells influences bone marrow stromal cell phenotype and compromises bone tissue quality throughout development and during bone response to voluntary wheel running. After successfully generating a LINC disruption mouse model under the Prrx1-Cre driver, we found no change in osteogenic and adipogenic differentiation potential in PDLD cells, nor did we find any bone architecture changes in PDLD mice at an 8-week baseline. While histology outcomes found that exercise-induced osteoid volume and surface was abolished in PDLD animals, there were no remarkable differences in bone microarchitecture or mechanical properties between PDLD mice subjected to a six-week exercise intervention compared to LINC intact animals or sedentary controls.

Studies involving LINC complex disruption via KASH overexpression *in vitro* show decreases osteogenic activity in human bone marrow mesenchymal stem cells by reducing Runx2 via increased histone acetylation [47]. We recently reported that LINC disruption also decreases adipogenic potential *in vitro* [48]. Despite this evidence, our results indicate no change in adipogenic and osteogenic potential of these cells when extracted from bone marrow. To exclude the possibility of these cells not surviving the marrow extraction we first confirmed that eGFP-KASH expressing LINC-deficient cells were present in our cultures after marrow extraction and next we ensured that LINC-deficient cells were participating in osteogenic nodules (Fig. S2). Finally, to test possible the of acute versus chronic effects of KASH protein expression, we utilized transient transfection with a previously published dominant negative form of the nesprin-1 KASH domain (DNK) [30, 31]. The DNK plasmid utilized contains a mutated form of the nesprin-1 KASH domain where the eGFP-KASH2 fusion protein contains a mutated form of the nesprin-2 KASH domain. While KASH inserts come from different nesprin isoforms the KASH domain is highly conserved between nesprin isoforms and results should be comparable. [49] RNA-seq and qPCR results from transient DNK transfection show decreased osteogenic gene transcripts *in vitro* when compared to empty mCherry plasmid transfected controls (Fig.S3). While it is not clear why we did not see any effect on cell phenotypes from animals, pathway analysis in DNK expressing cells show decreased levels of collagen deposition related genes (Table S1), which support the decreased osteoid volume and surface measures in PDLD animals (Fig.8). However, beyond osteoid measure we did not see any other skeletal effects in animals, therefore, it is possible bone cells in our animals may have adapted to this chronic expression of eGFP-KASH fusion protein and respond differently when compared to acute expression of Nesirin-1 KASH domain.

*Prrx1* is a well-established marker for early limb bud development [8, 9], and this study provides valuable insights into the impact of LINC disruption across various developmental stages, from embryogenesis through adolescence into adulthood. Interestingly, our findings suggest that this disruption has no significant influence on the differential potential of bone marrow cells or whole bone structural or mechanical properties. One possible explanation for this observation is that cells with LINC disruption might be selectively excluded from the population of bone-forming cells, possibly directed towards apoptotic pathways. Previous reports indicated a function of nesprin-2 in apoptotic cells where nesprin-2 binds to Bax and the binding increases in apoptotic cells[50] which may support this phenomenon. However, it’s worth noting that we do detect *Prx1-Cre* activity in adult bone marrow cell aspirates, indicating that these cells are still present in the adult skeletal system. Yet, their precise function and role in bone biology remain unclear. Some studies propose that cells originating from the Prx1 lineage may not be actively involved in the homeostatic mechanisms or bone remodelling [29]. This raises questions about the specific contributions of these cells within the intricate landscape of bone development and maintenance, prompting further investigation into their functions and potential implications for bone health.

Voluntary wheel running studies also with a six-week intervention but using older (16 week) female mice found increased trabecular quality and quantity post-exercise in animals on a regular diet [51] [52]. While bones of male mice also shown to respond to voluntary wheel running [53] [54] [55], other studies using male mice reported minimal effects of exercise on bone after 10 or 21 weeks of voluntary wheel running [56, 57]. Compared to reported daily running metrics in these studies (ranging from 4-10 km/day), our mice ran less on average (2.1 km/day). The reason for this is unknown, but may have affected our outcomes. Furthermore, our mice were 8-weeks old when placed in the exercise intervention, which is before the 12-week reported age of skeletal maturity [56, 58]. Therefore, it is possible that mechanical challenge provided by voluntary wheel running did not affect the ongoing bone growth in these mice and thus were not able to alter osteoblast numbers (Fig.8). Further indicated in Figure 8 however, we found however that voluntary running exercise significantly increases osteoid volume (OV/BV) and osteoid surface (OS/BS) in control animals indicate that exercise increases organic portion of the bone matrix that forms prior to the maturation after 6wk of voluntary running at 16wk of age. In PDLD mice however this response to voluntary running exercise was completely abolished suggesting Prrx1+ cells may play a role in immature bone deposition. Further studies using different modalities of LINC disruption (i.e. SUN & Nesprin depletion models) or the use of different mesenchymal/osteochondroprogenitor cre drivers (i.e. Dermo1 or Sox-9) would shed further light on the impact of impaired mechanotransduction in skeletal stem/progenitor cells.

## Acknowledgements

This study was supported by AG059923, AR075803, P20GM109095, NSF1929188 and NSF 2025505 to GU and AR074473 to WRT. The author(s) declare no competing interests financial or otherwise.

## Author Contributions

All authors have read and approved the final submitted manuscript.

Scott Birks: concept/design, data analysis/interpretation, manuscript writing.

Sean Howard: data analysis/interpretation, manuscript writing.

Caroline O’Rourke: data analysis/interpretation, manuscript writing.

Matthew Goelzer: data analysis/interpretation, manuscript writing.

William R Thompson: data analysis/interpretation, financial support, manuscript writing.

Anthony Lau: data analysis/interpretation, financial support, manuscript writing.

Gunes Uzer: concept/design, data analysis/interpretation, financial support, manuscript writing.

## Supplementary Figures

**Figure S1.**
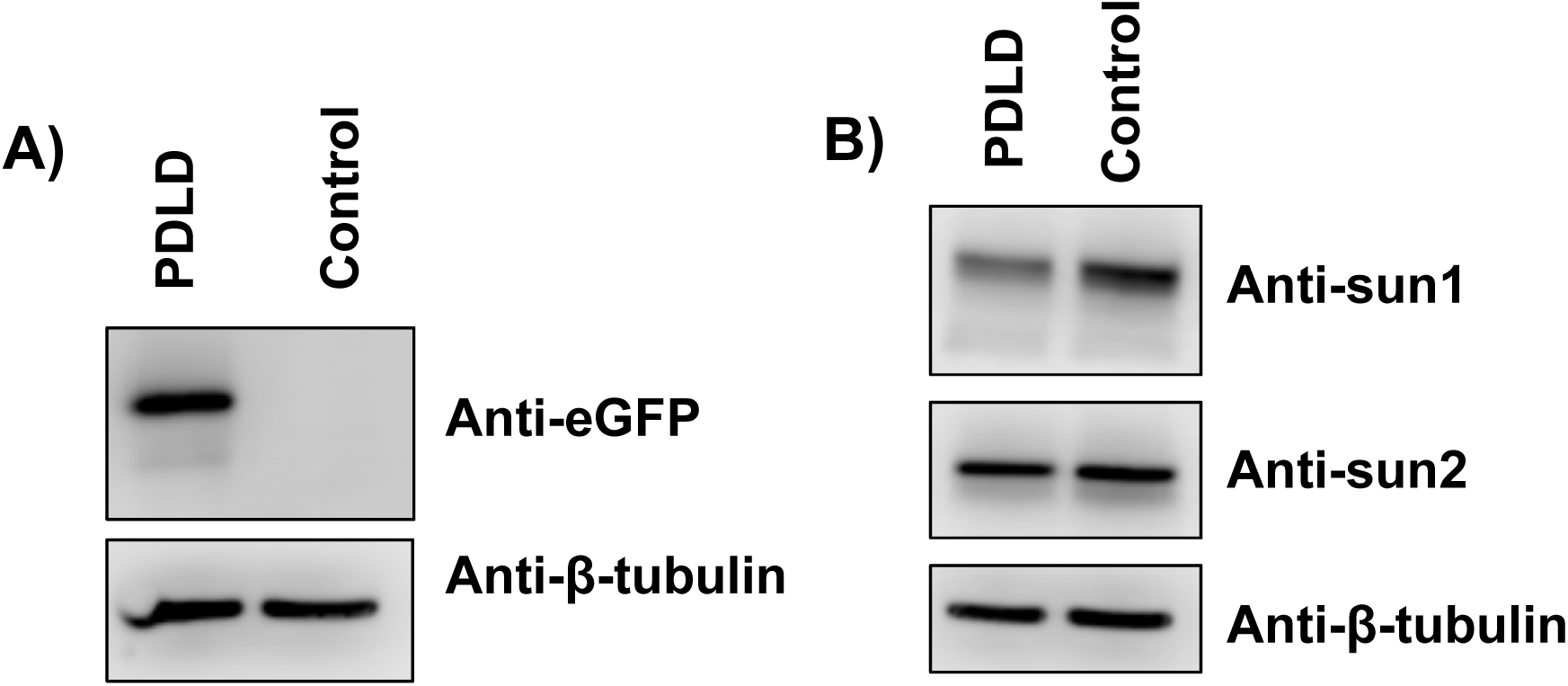
Western blot against eGFP showing **(A)** the presence of eGFP in bone marrow cells of PDLD mice and not in control mice. Anti-B tublin was used as a positive control. **(B)** indicates decreased levels of sun1 and sun2 in PDLD mice.

**Figure S2.**
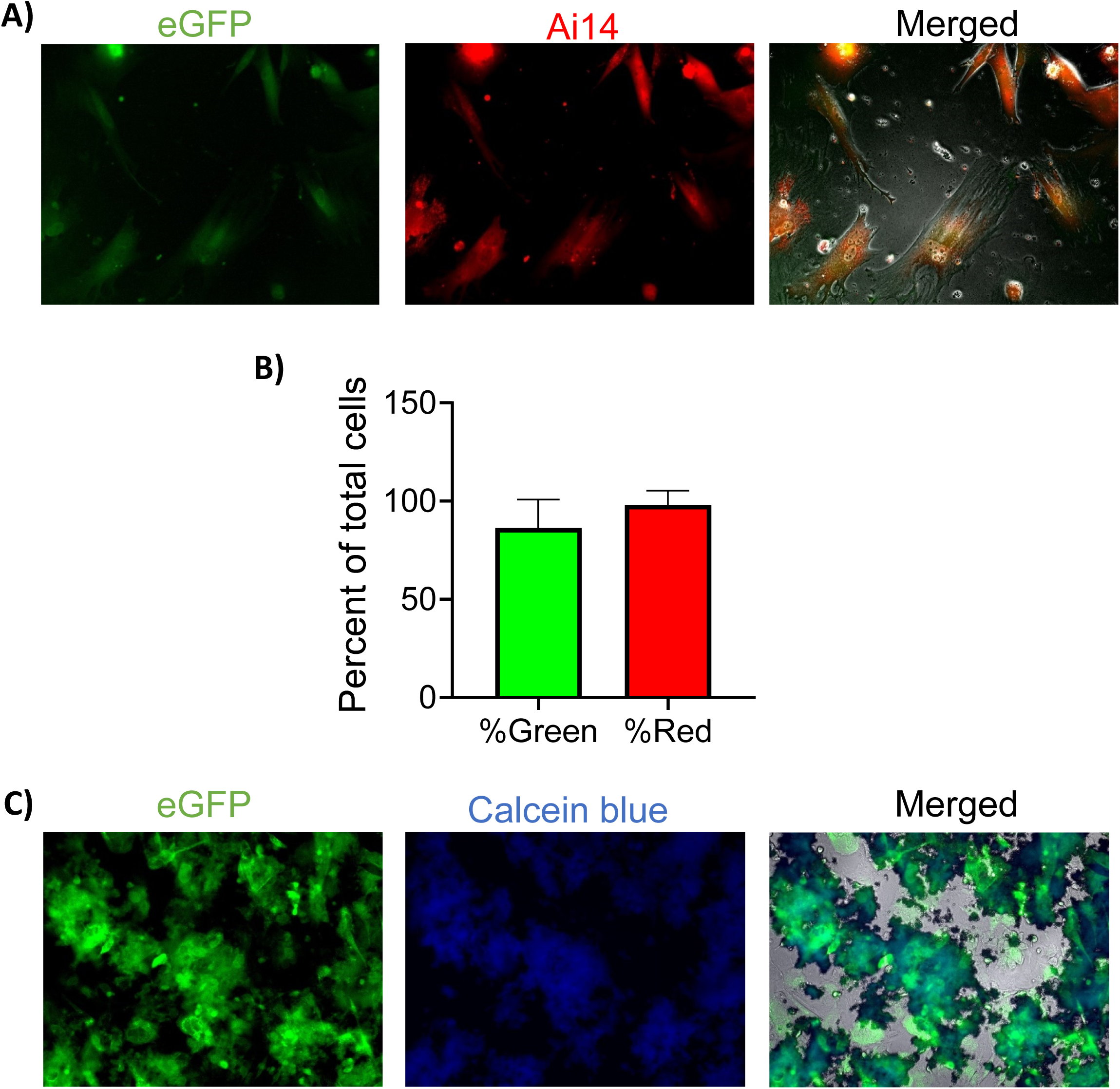
eGFP-KASH expressing cells in primary cell culture. **A)** Representative images of bone marrow aspirated from 12-week female Prx-Cre(+)/Ai14(+)/KASH(+) Number of cells per field expressing the GFP-KASH construct was determined by fluorescent microscopy in the control cells of Prx lineage were determined by tdTomato expression in both the control All counts were done utilizing FIJI. **B)** Graph of each cell percentages to total, error bars represent standard deviation. **C) KASH-GFP expressing cells contribute to the formation of osteogenic nodules.** Images were taken at 10X using a) FITC filter showing eGFP expression b), DAPI filter showing calcein blue fluorescence c) an overlay to qualitatively show that the green KASH-GFP expressing cells were located in the blue Calcein Blue stained nodule region

**Figure S3.**
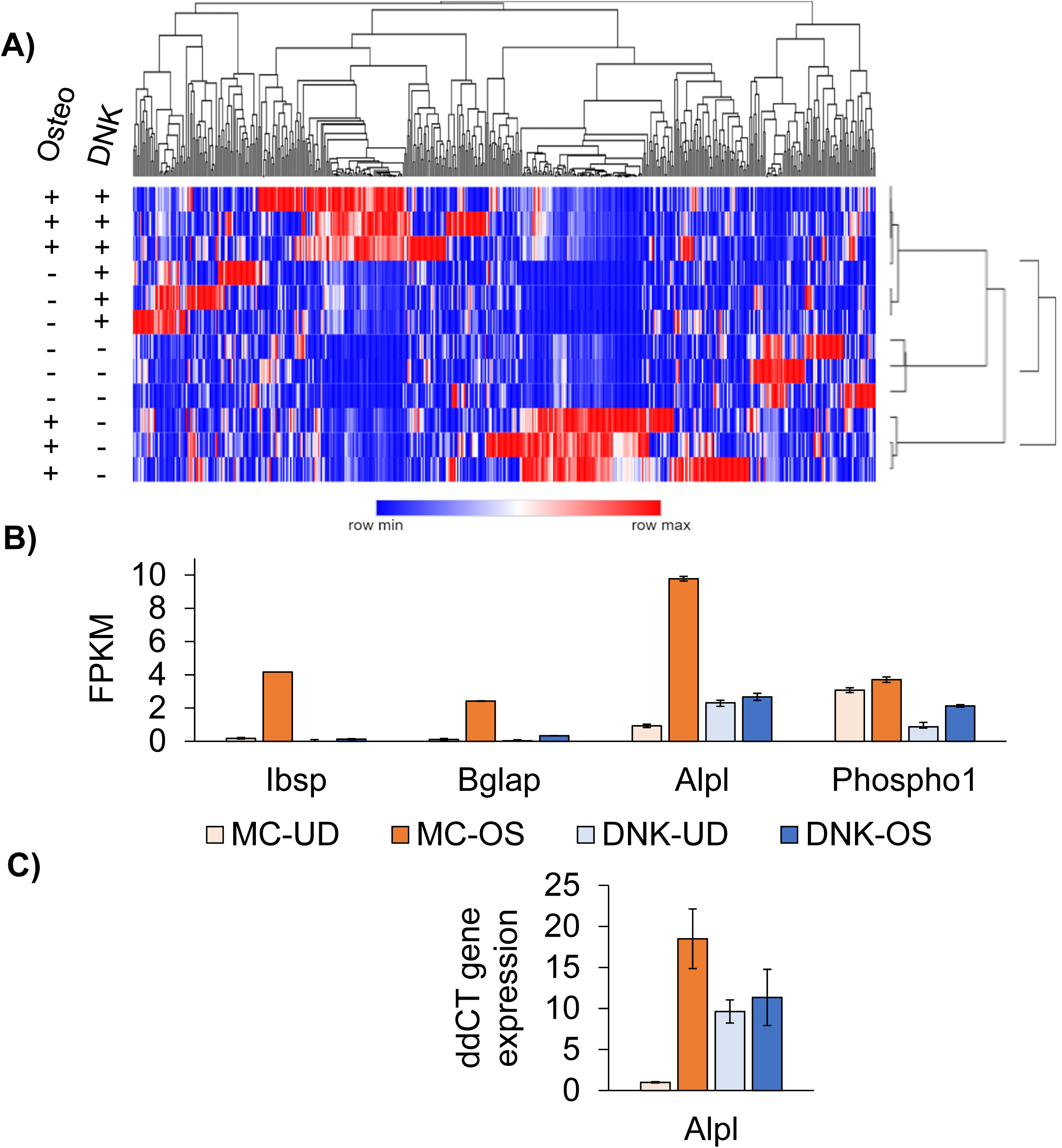
RNA-seq Analysis of DNK and MC Expressing MSCs During Osteogenesis: (**A**) Heatmap of FPKM values for mCherry tagged dominant negative KASH (DNK) and empty mCherry (MC) plasmids expressing MSCs during 7 days of osteogenesis all shown values were p<0.05. (**B**) FPKM values of osteogenesis genes of Ibsp, Bglap, Alpl, and Phospho1 in MSCs expressing DNK or MC plasmid during 7-day osteogenesis (**C**) qPCR quantification of same samples using Alpl primer (Forward: aacccagacacaagcattcc, Reverse: gcctttgaggtttttggtca). Both undifferentiated controls (UD) and osteogenic media (OS) was shown.

**Figure S4.**
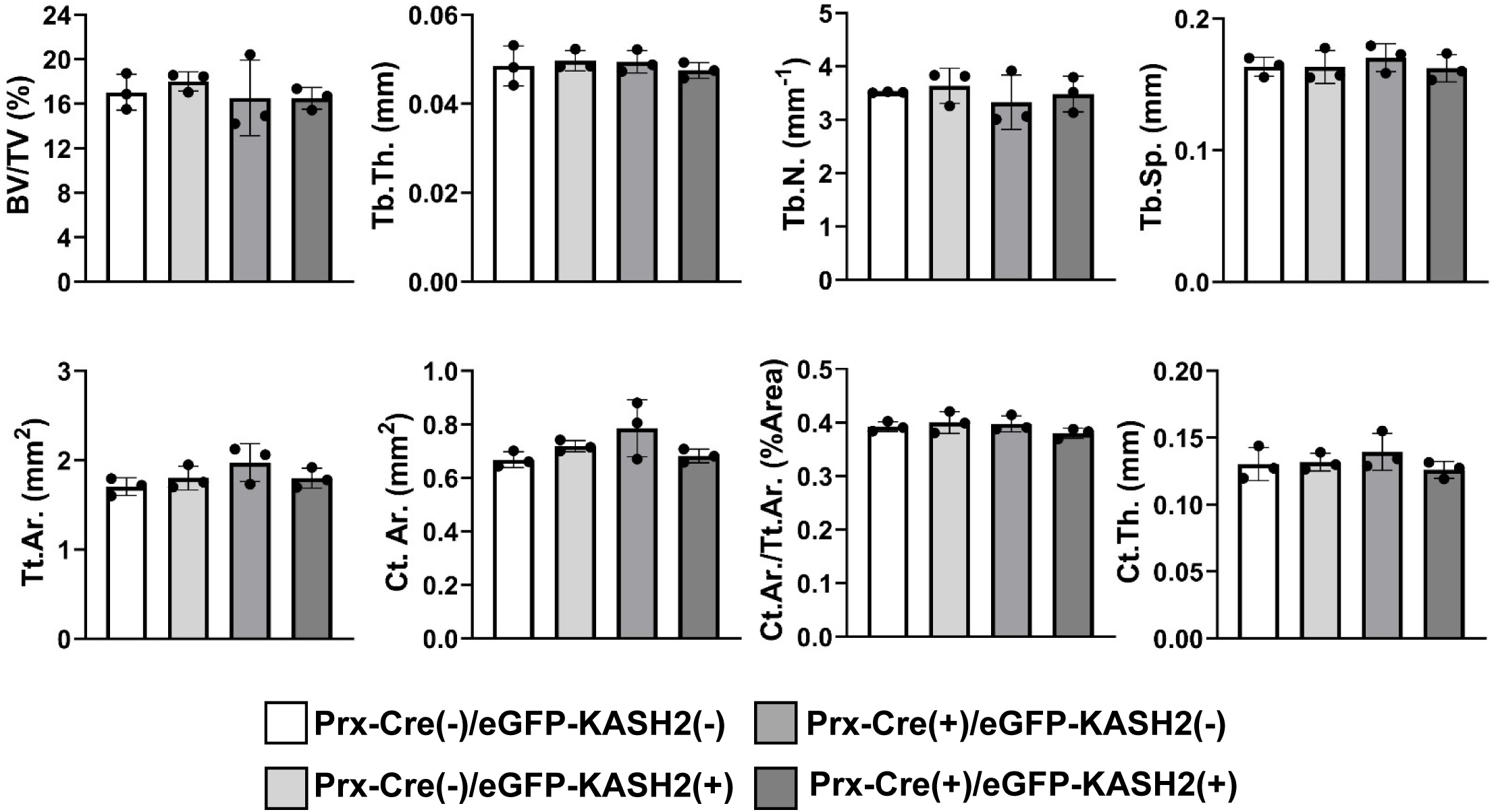
Bone microarchitecture for all four possible genotypes at 8-weeks. (n=3/grp): Prx-Cre(+)/EGFP-KASH2(+), Prx-Cre(+)/EGFP-KASH2(-), Prx-Cre(-)/EGFP-KASH2(+), Prx-Cre(-)/EGFP-KASH2(-). There was no significant difference found between genotypes. Significance determined via ordinary one-way ANOVA.

**Figure S5.**
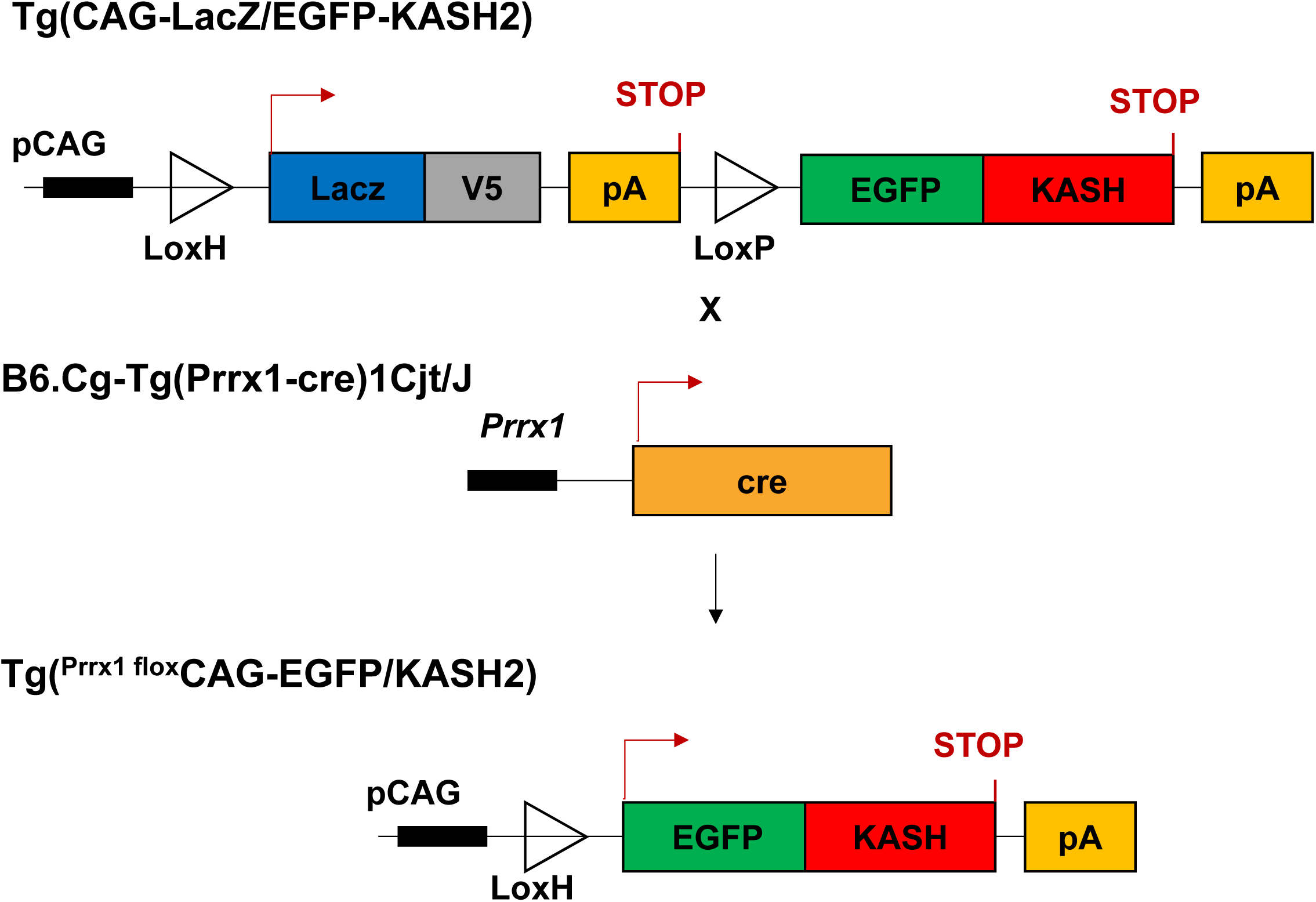
Breeding scheme for the creation of PDLD mice. Hemizygous Prx-Cre mice were crossed with floxed KASH2 mice to generate Prrx1-driven LINC disrupted (PDLD) murine model. LINC disruption mechanism described by Razafsky and Hodzic. Genotyping was performed on tissue biopsies via real-time PCR probing for LacZ and Cre (Transnetyx, Cordova, TN) to determine experimental animals and controls. Cre(+)/LacZ(+) animals were considered experimental and all other genotypes were considered controls.

**Figure S6.**
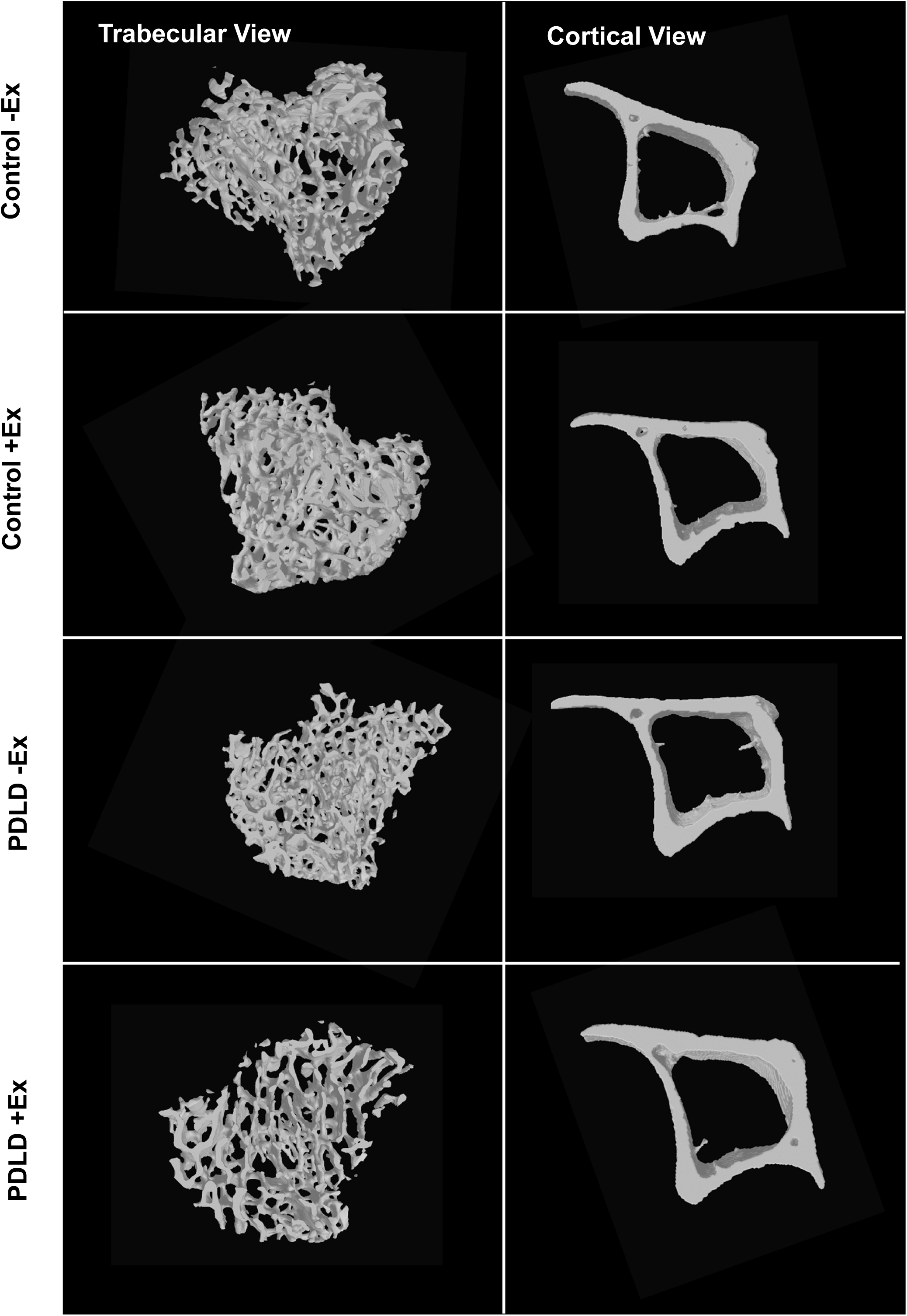
Representative micro-CT images for trabecular and cortical architecture of control and PDLD animals with and without exercise.

